# Inducible overexpression of zebrafish *microRNA-722* suppresses chemotaxis of human neutrophil like cells

**DOI:** 10.1101/578583

**Authors:** Alan Y. Hsu, Sheng Liu, Kent A. Brasseale, Ramizah Syahirah, Jun Wan, Qing Deng

## Abstract

Neutrophil migration is essential for battling against infections but also drives chronic inflammation. Since primary neutrophils are terminally differentiated and not genetically tractable, leukemia cells such as HL-60 are differentiated into neutrophil-like cells to study mechanisms underlying neutrophil migration. However, constitutive overexpression or inhibition in this cell line does not allow the characterization of the genes that affect the differentiation process. Here we apply the tet-on system to induce the expression of zebrafish *miR-722* in differentiated HL-60. Overexpression of *miR-722* reduced the mRNA level of genes in the chemotaxis and inflammation pathways, including *RAC2*. Consistently, cell migration is significantly inhibited upon induced *miR-722* overexpression. Together, *miR-722* is an evolutionarily conserved suppressor for neutrophil migration and signaling in zebrafish and human.

## Introduction

The neutrophil is the most abundant white blood cell in the circulation and a significant regulator of inflammation. While essential for battling against pathogens, neutrophil activation drives immunopathology in numerous human diseases, including organ transplantation, sepsis, rheumatoid arthritis, diabetes, neurodegenerative disease and cancer (1–3), although the link of some diseases to the innate immune system is not intuitive. Besides killing pathogens, they communicate with other cells to shape the inflammatory response. For example, neutrophils initiate inflammation by scanning platelets (4), migrating away from the initial activation site to disseminate inflammation to the lung (5), priming macrophages by providing DNA in the forms of extracellular traps (6) and can directly present antigens to activate T cells (7,8).

Manipulating neutrophil migration and activation is implicated in managing chronic inflammation (3,9). The current challenge in the field is that primary neutrophils are terminally differentiated with a very short life span ex vivo, which excludes the feasibility of genetic manipulation for functional characterization. To model neutrophils, human promyelocytic leukemia HL-60 cells (10) and NB4 cells (11) are differentiated in culture for 5-7 days into neutrophils-like cells. Gene transduction approaches using either the lenti- or retro-virus are successful in these cell lines, providing a genetically tractable system. However, due to the cell differentiation process, extensive characterization is required to separate the effect of the target genes on cell differentiation and mature cell function. Inducible expression using tet-on (gene expression activated by doxycycline) is widely applied in cancer research and other cell lines. However this technique has not been applied to neutrophil precursor cells.

MicroRNAs (MiRNA) are evolutionarily conserved, non-coding RNAs of ~22 nucleotides that post-transcriptionally regulate gene expression (12). miRNAs are master regulators that can simultaneously suppress hundreds of genes and regulate numerous cellular processes and human diseases. miRNA profiles are distinct in human peripheral blood neutrophils (13–15) and activated tissue infiltrating neutrophils (16), suggesting that they are regulated by the inflammation process or tissue environment. On the other hand, only a few miRNAs are functionally characterized in neutrophils (17). In HL-60 cells, miRNA expression changes during cell differentiation (18–20) and after radiation (21) or resveratrol treatment (22). Indeed, multiple miRNAs regulate HL-60 growth, differentiation and survival in vitro (20,23–31). On the other hand, reports on how miRNAs regulate differentiated cell function such as migration is scarce. Introduction of synthetic miR-155 and miR-34 mimics into differentiated HL-60 suppressed cell migration but not differentiation (32), whereas depleting miR-155, on the other hand, induced cell differentiation and apoptosis (33).

In zebrafish, we have identified a miRNA, *miR-722*, that when overexpressed in neutrophils, reduces neutrophil chemotaxis and protects the organism from both sterile and non-sterile inflammatory assaults (34). A hematopoietic specific isoform of the small GTPase, *rac*2, was identified as a direct target of *miR-722*. Here, we used the tet-on technique to induce the overexpression of *miR-722* in differentiated HL-60 cells and uncovered an evolutionarily conserved function of *miR-722* in neutrophils.

## Results

### Establishing an inducible gene expression system in HL-60

To establish a platform that allows inducible gene expression in HL-60 cells, a lentiviral backbone initially constructed by Dr. David Root (Addgene plasmid # 41395) was selected. It contains all elements to enhance viral production and integration, including a Psi packaging element to facilitate pseudovirus production, REE and WPRE nuclear exporting elements to enhance integration in the host genome. The puromycin resistant gene and the reverse tetracycline-controlled transactivator is constantly expressed under the control of the PGK promoter. Addition of tetracycline or its close derivative doxycycline induces the binding of the transactivator to the tet responsive element, activating the target gene expression in a dose dependent manner (Figure 1A). The microRNA hairpin is cloned into the intron of a fluorescent reporter gene, *Dendra2*. To ensure an effective selection of cells with integration, we measured cell proliferation in the presence of increased doses of puromycin. HL-60 cells are killed 3 days post treatment under all tested conditions, and we selected the condition of 1 µg/ml puromycin treatment for at least 7 days. (Figure 1B). We then tested whether differentiated HL-60 cells (dHL-60) can tolerate doxycycline treatment. Cells 4 days post differentiation were treated with increased doses of doxycycline, and cytotoxicity was observed at concentrations higher than 1 µg/ml (Figure 1C).

**Figure 1.**
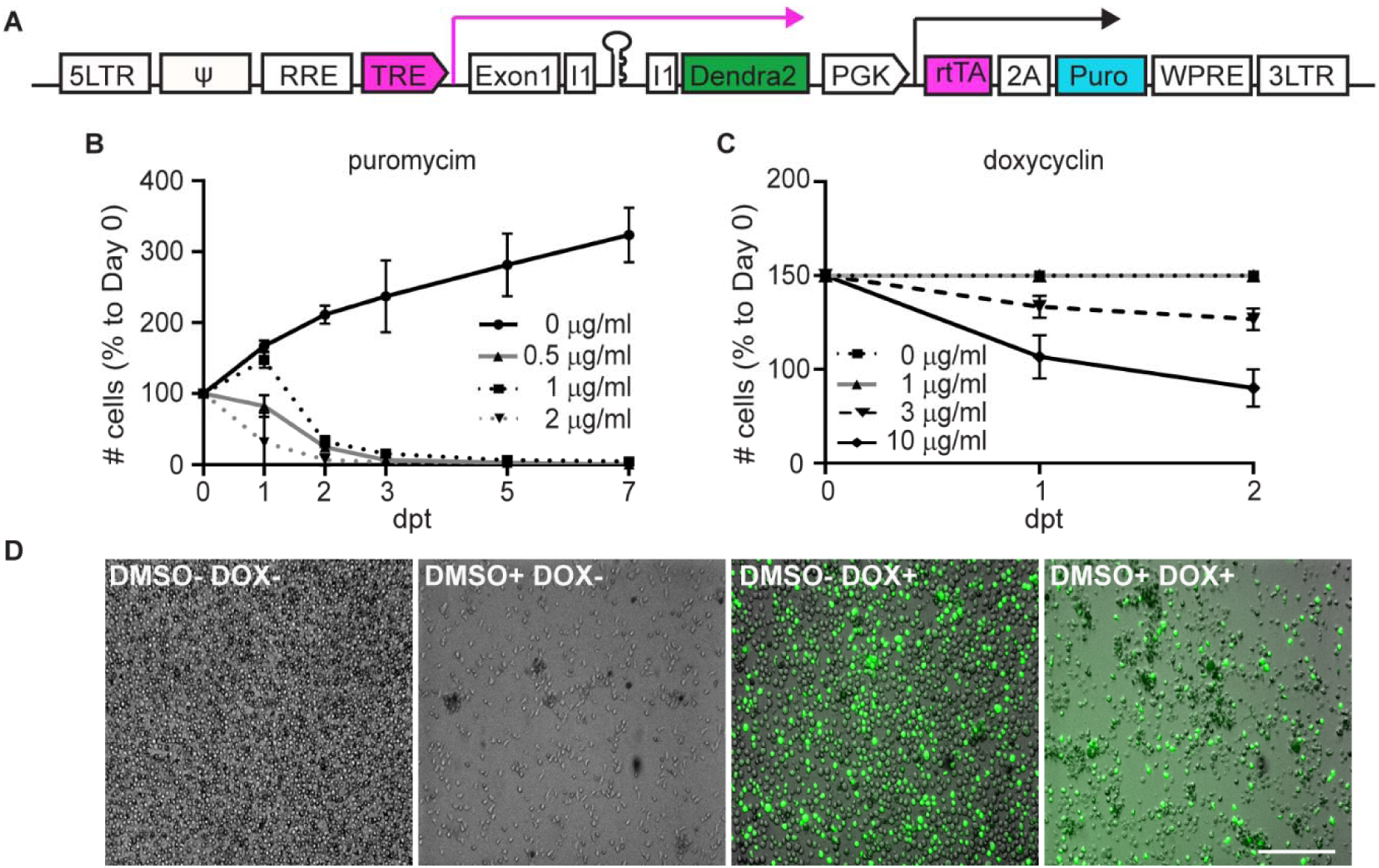
Inducible expression of a miRNA together with a reporter gene in HL-60 cells. (A) Schematics of the plix 4.03 vector modified for the inducible expression of a microRNA and a reporter. The miRNA hairpin resides in the intron of the Dendra2 reporter (green) under the control of a doxycycline response element (TRE, magenta and magenta arrow). The puromycin resistant cassette (Puro, cyan) linked with the reverse tetracycline-controlled transactivator (rtTA, magenta) was driven by a constitutively active PGK promoter (black arrow). Flanking elements including the Psi packaging element, REE and WPRE nuclear exporting elements enhance integration in the host genome. (B) Proliferation of HL-60 cells in the presence of the indicated doses of puromycin. Cell numbers were normalized to day 0; dpt: days post treatment. (C) Survival of differentiated HL-60 cells in the presence of the indicated doses of doxycycline starting 4 day post differentiation. Cell numbers were normalized to day 0; dpt, days post treatment. (B, C) Results are averaged of three independent experiments and shown as mean ± s.d.. (D) Representative images of a stable line of HL-60 cells transduced with the construct depicted in (A) with/without differentiation (± DMSO) or induction (± DOX). Scale bar: 200 µm.

### Inducible expression of *miR-722* in HL-60

We then generated stable HL-60 lines in which the *miR-722* or a vector control expression is regulated by the tet responsive element. For simplicity, we refer to them as *miR-722* and control lines. After induction, we observed 50-60 % of cells expressing detectable levels of the Dendra2 reporter with or without DMSO induced differentiation (Figure 1D). The experimental flow thus is finalized as depicted in Figure 2A. To confirm that *miR-722* expression is indeed induced, we quantified the level of the precursor transcript and the mature microRNA. Since *miR-722* is absent in human and HL-60 cells, we detected 10,000 fold increase of the mature *miR-722* level in the *miR-722* line, but not in the control line (Figure 2B). General microRNA biogenesis was not affected, since the level of two other mature microRNA, *MIR-223* and *LET7* in dHL-60 was not affected. Additionally, cell viability or differentiation (as measured by surface expression of annexin V binding and CD11b expression) was not affected by overexpression of *miR-722* in differentiated HL-60 cells (Figure 2C, D).

**Figure 2.**
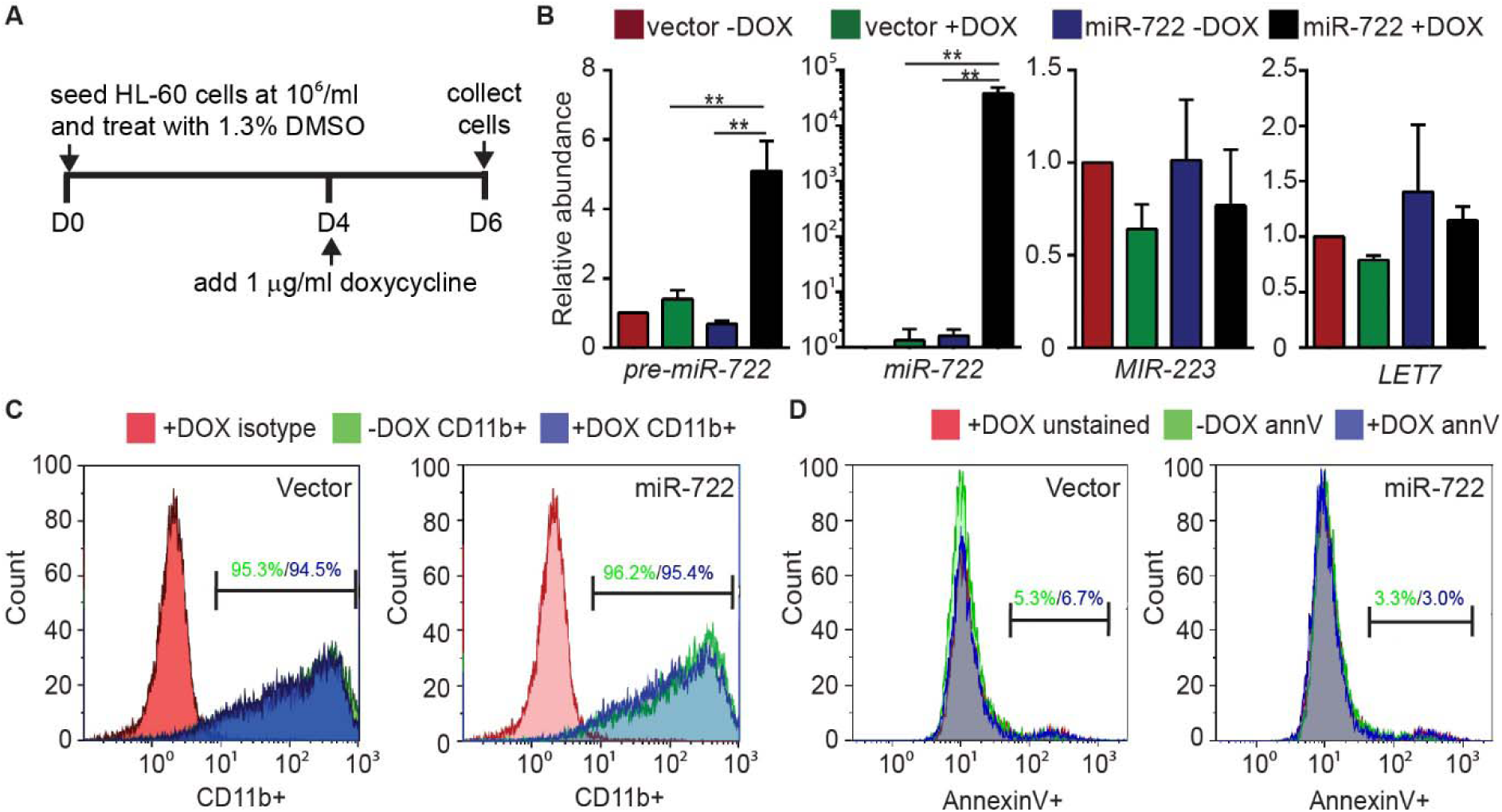
Specific induction of *miR-722* does not affect HL-60 differentiation or survival. (A) Schematics of the experimental flow for the inducible expression of a microRNA and a reporter. The stable HL-60 cell lines over expressing *miR-722* or the vector control were differentiated with 1.3 % DMSO for 6 days and treated with 1 µg/ml doxycycline from day 4 or left untreated. (B) Levels of *pre-miR-722*, *miR-722*, *MIR-223*, and *LET7* in vector or miR-722 expressing dHL-60 lines ± doxycycline (DOX). Messenger RNAs were normalized to *GAPDH* and miRNAs were normalized to *U6*. Results are presented as mean ± s.d. from three independent experiments. **, p < 0.01, Mann–Whitney test. (C, D) Histogram of CD11b expression (C) and Annexin V staining (D) of vector and *miR-722* expressing dHL-60 with (blue) or without (green) DOX. Positive cells are gated based on the isotype control and percentages of positive cells are labeled in corresponding color. Images are representative of three independent experiments.

### *miR-722* suppresses cell migration and signaling related genes in HL-60

To characterize the signaling pathways that are regulated by *miR-722*, we performed RNAseq of the *miR-722* and the control lines. Since lentiviral mediated DNA integration is random, although with some preferences (35,36), we compared the gene expression changes in both lines ± Dox to minimize the artifact associated with insertion sites. Induction of *miR-722* in dHL-60 resulted in significant dis-regulation of transcripts (FDR-adjusted p-value < 0.01 & FC < -log_2_1.8, see Methods for details) as shown in Figure 3A and Supplementary Dataset 1. There were much more down-regulated differentially expressed genes (DEGs) (n = 1943) identified than up-regulated ones (n = 632), which is consistent with the function of microRNAs as suppressors of gene expression. Among the down-regulated transcripts, the human orthologues of predicted *miR-722* targets in zebrafish were significantly enriched (p = 0.003) (Figure 3B), while *miR-722* targets were depleted in up-regulated gene group (p = 0.031). The results indicate an evolutionary conservation of *miR-722* targets. Under the same cutoffs used in comparison in the *miR-722* line, no significant dysregulation was identified in the control line ± Dox, where only 15 transcripts had p < 0.01 without multiple-test correction (Supplementary Figure 1 and Supplementary Dataset 2). Moreover, fold changes of genes in the control line were presented in a symmetric way between up- and down-regulation. The different patterns between *miR-722* and vector suggests that the effect of *miR-722* was not due to the reporter gene expression alone. Via enrichment analysis on Gene Ontology (GO) and KEGG pathways, we found that the down regulated transcripts were significantly over-represented in pathways regulating cell migration, chemotaxis, signaling and inflammation (Figure 3C, D, E, F and Supplementary Dataset 3). To validate the RNAseq results, we quantified the relative expression level of several transcripts in the chemotaxis pathway and indeed all were significantly downregulated upon *miR-722* induction, but not the reporter gene alone (Figure 4A). Among those, the chemokine receptor encoding gene, *Fpr1*, harbors *miR-722* binding sites in its 3′UTR and is a potential direct target. Since *rac2* is a direct target of *miR-722* in zebrafish, we tested whether human *RAC2* is also a *miR-722* target. We detected significant down regulation of *RAC2* at both the transcript and protein levels upon *miR-722* overexpression (Figure 4A, B). Indeed, a 7mer-m8 site (complementarity at nt 2-7 and mismatch at nt8) was present in the *RAC2* 3′UTR (Figure 4C). In a luciferase reporter assay, specific suppression by *miR-722* was observed with the wild-type *RAC2* 3′UTR, but not the mutated 3′UTR lacking the seed binding site (Figure 4D, E), confirming that *RAC2* is a direct target of *miR-722* in human cells.

**Figure 3.**
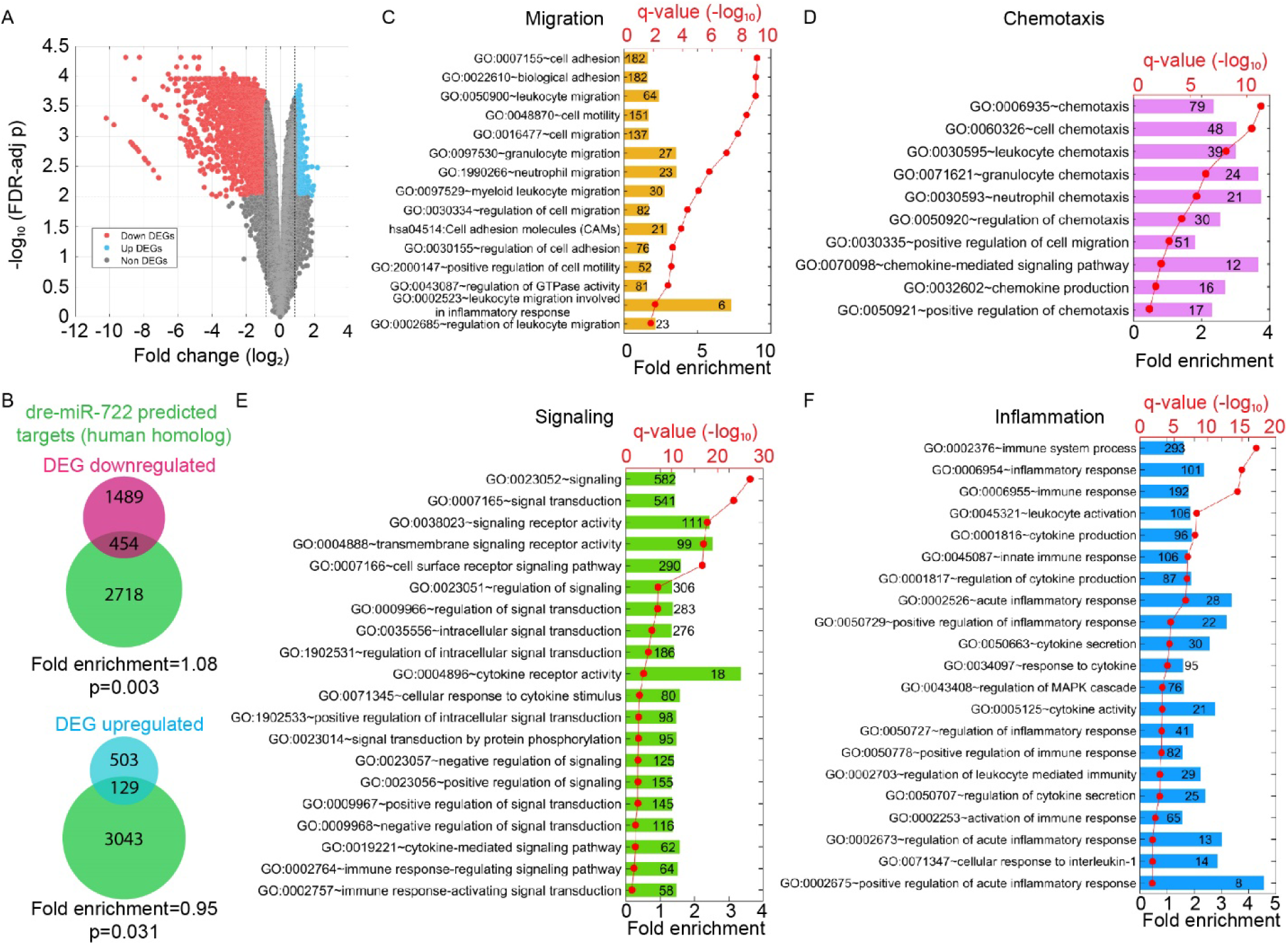
*miR-722* overexpression suppresses the expression of cell signaling and migration related genes in dHL-60. (A) Volcano blot of DEGs with significant changes in expression upon doxycycline induced *miR-722* expression. Red: down-regulated differentially expressed genes (DEGs); cyan: up-regulated DEGs. (B) Significant enrichment of human homologues of *miR-722* predicted target genes in the down-regulated DEGs. P < 0.003, hypergeometric distribution. (C, D, E, F) Clusters of the genes in the pathways significantly altered upon *miR-722* overexpression. Number of the genes with significant expression changes in each pathway are labeled in each column. q-values, FDR-adjusted p-values as described in the Methods section.

**Figure 4.**
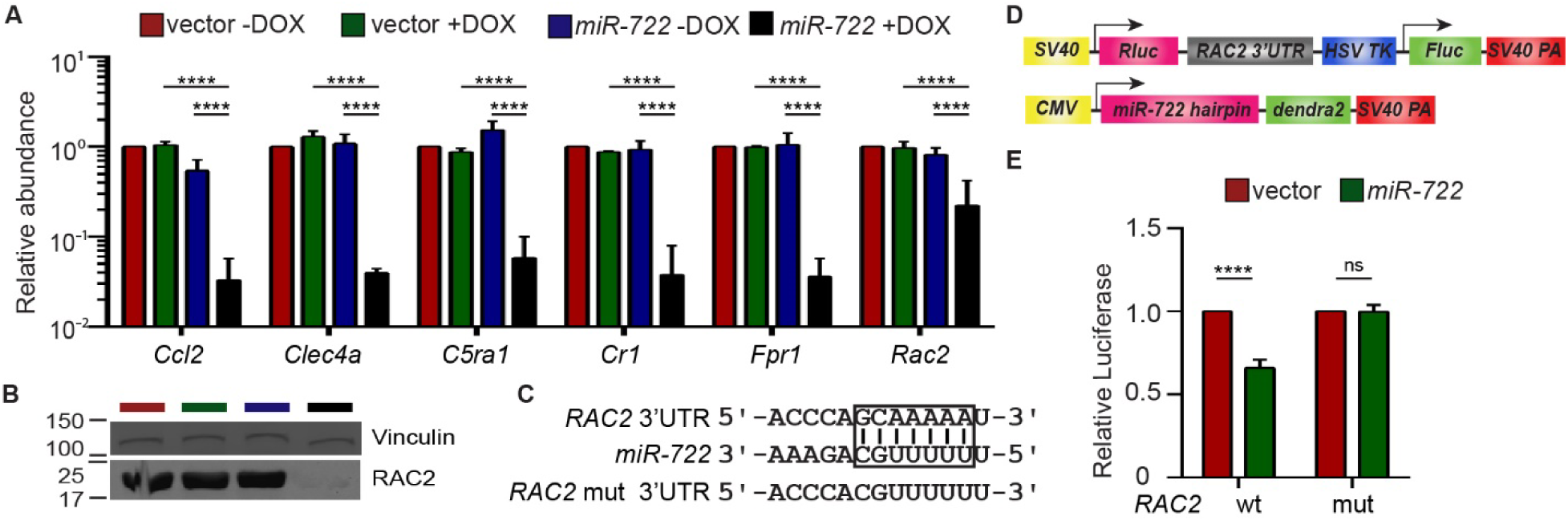
*RAC2* is a direct target of *miR-722*. (A) Relative abundance of mRNA levels of indicated genes in vector or *miR-722* expressing dHL-60 ± DOX. Results are presented as mean ± s.d. from three independent experiments. ****, p < 0.0001, Sidak’s multiple comparisons test. (B) Immunoblot of RAC2 in vector or *miR-722* expressing dHL-60 cells ± DOX. Vinculin is used as a loading control. The blot was cut in half and probed with two different primary antibodies. Cropped images from the full length blot (Supplementary Figure 2) are shown. (C) Alignment of *miR-722* seed sequence and the 7mer-m8 seed binding site in the *RAC*2 3′UTR which is then mutated for the reporter assay in (E). (D) Schematics of the constructs used for the luciferase reporter assay. Wild-type or mutated *RAC2* 3′UTR were cloned downstream of a renilla luciferase gene. A firefly luciferase gene on the same plasmid was used as a normalization control. *mir-722* or vector hairpin was cloned into a mammalian expression plasmid for expression. (E) Selective suppression of renilla luciferase activity by *miR-722* through binding to the seed sequence in *RAC*2 3′UTR. Results are presented as means ± s.d. from three independent experiments. ****, p < 0.0001, Mann–Whitney test.

### *miR-722* suppresses chemotaxis in HL-60

*RAC2* is essential for actin polymerization and migration in neutrophils (37). We then measured chemotaxis of the *miR-722* line using two independent assays. First, cells were allowed to migrate to the chemoattractant fMLP in a chemotaxis slide. *miR-722* overexpressing cells displayed a significant defect in velocity but not in directionality, compared with the vector control or the *miR-722* expressing line without doxycycline mediated induction (Figure 5A-C, Supplementary Movie 1, 2). Consistently, *miR-722* overexpression suppressed cell transwell migration (Figure 5D), supporting the functional inhibition of the *RAC2* gene in dHL-60s by *miR-722*.

**Figure 5.**
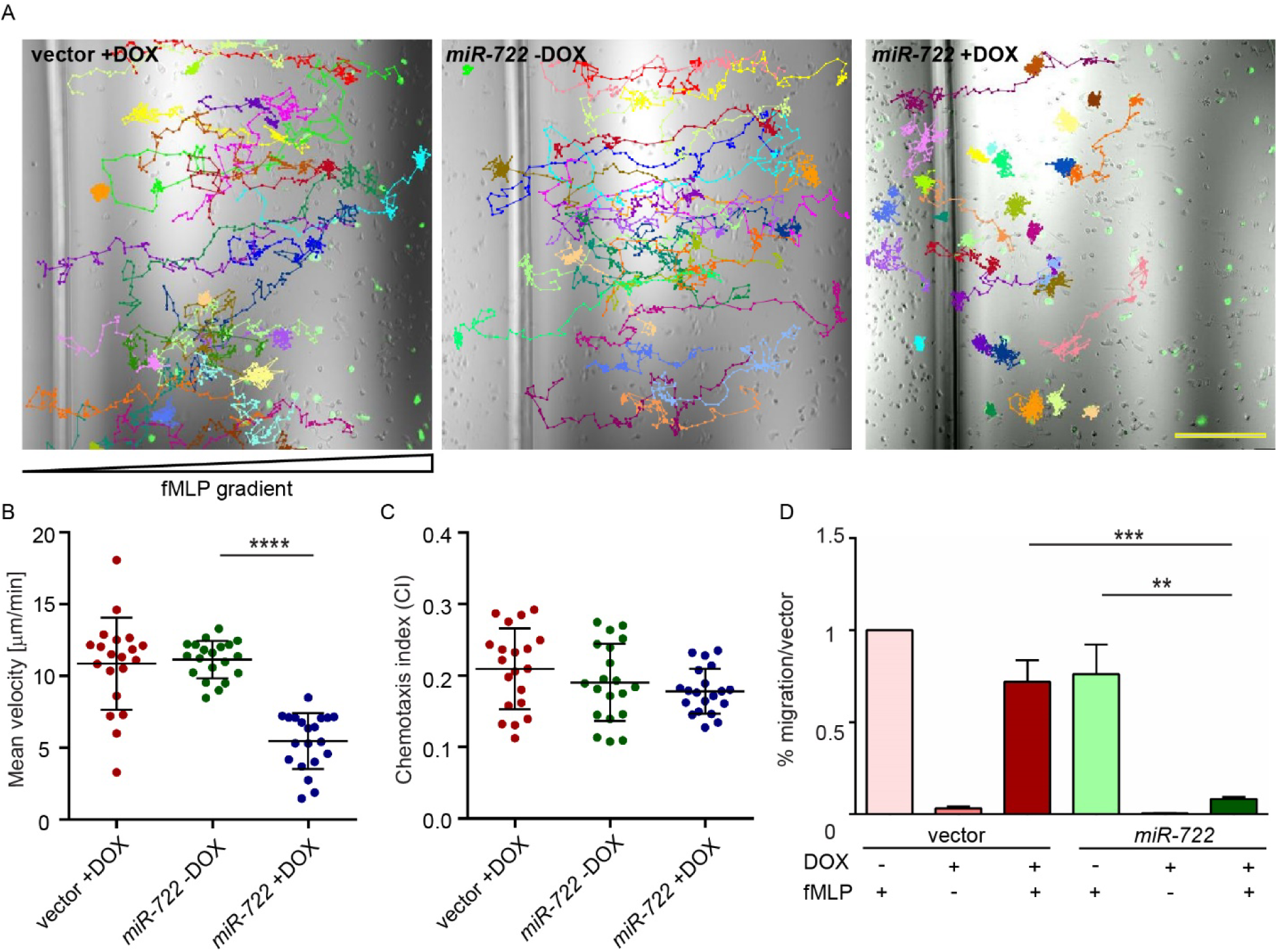
*MiR-722* overexpression suppresses HL-60 chemotaxis. (A) Representative images and tracks of dHL-60 cells expressing either vector (left) or *miR-722* with (right) and without (middle) doxycycline toward fMLP. Scale bar: 200 µm. (B, C) Quantification of velocity (B) and directionality (C) of dHL-60 cells during chemotaxis. Results are representative of three independent experiments. ****, p < 0.0001, Sidak’s multiple comparisons test. (D) Transwell migration of vector or *miR-722* expressing dHL-60 cells towards fMLP. Results are average of three independent experiments and presented as mean ± s.d.. **, p < 0.01; ***, p<0.001, Sidak’s multiple comparisons test.

## Discussion

Here we have applied the tet-on system in differentiated HL-60 (dHL-60) cells and uncovered an evolutionally conserved role of *miR-722* in suppressing neutrophil migration and signaling.

To our knowledge, it is the first application of the tet-on system in dHL-60 cells. Tet-on and tet-off system is favored in biomedical research because acute depletion or overexpression is more faithfully in reflecting the target gene function. The caveat of chronic depletion or overexpression is the possible compensatory expression of other off-target genes and altered overall cell physiology, which will complicate the result interpretation. An additional challenge with using dHL-60 cell model to study neutrophil function is the required differentiation process. It is essentially impossible to study genes that would affect cell differentiation without an inducible system. The dose of doxycycline dHL-60 could tolerate (1 µg/ml) is significantly lower than that in other cell lines (10 µg/ml). Although doxycycline is a broad spectrum antibiotic that inhibit bacterial protein synthesis, it also inhibits protein synthesis in eukaryotic mitochondria and thus can impair cell metabolism and cause cell death (38,39). The mechanism underlying this increased sensitivity is not clear but could be due to increased uptake and/or decreased efflux of doxycycline in dHL-60. In addition, doxycycline or its degradation products may be highly toxic to dHL-60s. We have optimized our protocol where the cytotoxicity of doxycycline is not observed. Cell proliferation, survival and differentiation were not affected by doxycycline treatment. We expect that our method is equally applicable to the inducible expression of other RNA species such as shRNAs in dHL-60, providing a platform for acute depletion of endogenous proteins to evaluate their biological significance in mature cell function without affecting the differentiation process.

Fundamental biological processes, such as cell proliferation and migration, are conserved through evolution. It is estimated that ~60 % of human protein-coding genes are under selective pressure to maintain miRNA seed binding sites in the 3′UTR which are conserved in vertebrates (40). It is not surprising that both human *RAC2* and zebrafish *Rac2* are direct *miR-722* targets. It is intriguing that *miR-722* expression in human HL-60 cells resulted in a down-regulation of genes that are significantly enriched for the orthologues of the predicted zebrafish targets. The significantly altered pathways provides explanation of the anti-inflammatory effect of this microRNA in zebrafish as its overexpression protected zebrafish against lethal inflammatory assault (34).

Rac2 plays a principal role in regulating the actin cytoskeleton, chemotaxis and signaling. Rac2 is required for neutrophil motility and chemotaxis (41), retention in the hematopoietic tissue (42), generation of super oxide ions during infections (43) and degranulation of primary granules (44). The therapeutic potential of Rac2 suppression in controlling inflammation has not been explored. Currently, a Rac2 specific inhibitor is not available and a complete Rac2 inhibition would result in primary immune deficiency and poor wound healing (45). Although the therapeutic potential of *miR-722* is not clear without further characterization, our work here has characterized the first microRNA that targets human *RAC2* and established *miR-722* as a evolutionarily conserved suppressor for neutrophil migration and signaling.

## Methods

### Generation of stable HL-60 cell lines

HEK-293 cells were cultured in DMEM supplemented with 10% FBS, 4.5 g/glucose and sodium bicarbonate. HL-60 cells were obtained from ATCC (CCL-240) and cultured using RPMI-1640 with HEPEs supplemented with 10% FBS with sodium bicarbonate. The lentiviral backbone pLIX_403 was a gift from David Root (Addgene plasmid # 41395). DNA sequence encoding microRNA-722 tagged with Dendra2 were amplified from (Addgene plasmid # 97163) with the following primers: pLIX-mir+: 5’-TGGAGAATTGGCTAGCGCCACCATGGATGAGGAAA TCGC-3’, pLIX-mir-: 5’-CATACGGATAACCGGTTACCACACCTGGCTGGGC-3’and cloned into pLIX_403 vector using the NheI/AgeI sites. Stable HL-60 cell lines were generated as described (46). Briefly, HEK-293 cells were transfected with pLIX_403, VSV-G and CMV8.2 using lipofectamine 3000 (ThermoFisher). The viral supernatant was harvested on day three and concentrated with Lenti-X concentrator (Clontech). HL-60 Cells were spin infected at 2000 g for two hours at 32 ºC and then selected and maintained in medium supplemented with 1 µg/ml puromycin. Live cells that excluded trypan blue stained cells were counted using a hemocytometer.

### Epifluorescence microscope imaging

Induction of the Dendra2 reporter expression was verified by imaging the green fluorescence using AXIO Zoom V16 microscope (Zeiss) at 110 % magnification. All images were acquired under the same condition within the linear range of the CCD detector.

### RT-qPCR

Total RNA was purified using MiRVANA miRNA purification kit (ThermoFisher). MicroRNAs were reverse transcribed with Universal cDNA Synthesis Kit II (Exiqon). MicroRNA RT-qPCR was performed with ExiLENT SYBR^®^ Green master mix (Exiqon) using LightCycler ^®^ 96 Real-Time PCR System (Roche Life Science). Primers used in this study are: miR-223-3p (205986), dre-miR-722 (2107521), hsa-let-7e-5p (205711) and human U6 (203907). Messenger RNAs were reverse transcribed with Transcriptor First Strand cDNA Synthesis Kit (Roche). RT-qPCR were performed with FastStart Essential DNA Green Master (Roche). Primers used are: pre722+: 5’-cggagtggaatttgaaacgttttggc-3’; pre722-:5′-cggagcgaaatctgaaacgtttctgc-3′; CCL2+: 5′-agtctctgccgcccttct-3′; CCL2-: 5′-gtgactggggcattgattg-3′; CLEC4A+: 5′-cccctcttgaggatatgtgc-3′; CLEC4A-: 5′-aggaaccaaacatggtggtaa-3′; C5AR1+: 5′-aagtaatgatacagagggatcttgtg-3′; C5AR1-: 5′-ttgcaagatgttgccattg-3′; CR1+: 5′-ctccccctcggtgtatttct-3′; CR1-: 5′-cctgtttcctggtactctaattgc-3′; FPR1+: 5′-tctctgctggtccccatatc-3′; FPR1-: 5′-ttgcaataactcacggattctg-3′; RAC2+: 5′-gatgcaggccatcaagtgt-3′; RAC2-: 5′-ctgatgagaaggcaggtcttg-3′; GAPDH+: 5′-ccccggtttctataaattgagc-3′; GAPDH-: 5′-cttccccatggtgtctgag-3′. The specificity of the primers were verified with a single peak in the melt-curve. The relative fold change with correction of the primer efficiencies was calculated following instructions provided by Real-time PCR Minor (http://ewindup.info/miner/data_submit.htm) and normalized to *U6* (miRNA) and *GAPDH* (mRNA) respectively.

### Flow Cytometry

Cells were stained with Alexa Fluor 647 conjugated CD11b (neutrophil differentiation marker, Biolegend, 301319, clone IV-M047) or the isotype control antibody (Biolegend, 400130, clone MOPC-21) and RUO AnnexinV (apoptosis marker, BD, 563973). Cells were stained in staining buffer (1%BSA 0.1% NaN_3_) and incubated on ice for 1 hour, washed three times with staining buffer and resuspended to suitable volume. Fluorescence intensity and collected using BD LSR Fortessa. Results were analyzed with Beckman Kaluza 2.1 software. The raw counts are provided in Supplementary Table 1.

### RNAseq

HL-60 cells were differentiated for 4 days in 1.3% DMSO (47), induced with 1µg/ml doxycycline or left un-induced for 2 additional days, and then pelleted into microtubes each containing 10^7^ cells. Total mRNA was extracted using RNeasy Plus Mini Kit (Qiagen #74104). RNAseq was performed at The Center for Medical Genomics at Indiana University School of Medicine. Samples were polyA enriched and sequenced with Illumina HiSeq 4000 with reads range from 37M to 44M. The RNA-seq raw data and processed data are in the process of being submitted to the Gene Expression Omnibus (GEO) and will be publicly available upon the acceptance of the manuscript.

### Bioinformatics analysis on RNA-seq data

The sequencing reads was mapped to the human genome using STAR v2.5 RNA-seq aligner (48) with the following parameter: “--outSAMmapqUnique 60”. Uniquely mapped sequencing reads were assigned to hg38 genes using featureCounts v1.6.2 (49) with the following parameters: “-s 2 –p –Q 10”. The genes were filtered if their count per million (CPM) of reads were less than 0.5 in more than 4 of the samples. The TMM (trimmed mean of M values) method was used for normalization on gene expression profile across all samples, followed by the differential expression analysis using edgeR v3.20.8 (50,51). The gene was determined as differentially expressed gene (DEG) if its p-value after FDR-adjusted multiple-test correction was less than 0.01 and its amplitude of fold change (FC) was larger than 1.8. We identified 1943 down-regulated DEGs and 632 up-regulated DEGs by comparing miR-722 ± Dox. However, no DEG was recognized for the control line based on the same cutoffs. The software package, DAVID Bioinformatics Resources 6.8 (52,53) was adopted for Gene Ontology (GO) and KEGG pathway enrichment analysis. The significant enriched GO terms or KEGG pathways were selected if their FDR-adjusted p-values (q-values) were less than 0.05 after multiple test correction, when we compared DEGs down-regulated by *miR-722* +Dox with all expressed genes tested by RNA-seq.

### Immunoblotting

HL-60 cells were differentiated for 4 days in 1.3% DMSO (47), induced with 1ug/ml Doxycycline or left un-induced for 2 additional days, and then pelleted into microtubes each containing 10^6^ cells. Cells were lysed with RIPA buffer (Thermo #89900) supplemented with proteinase inhibitor cocktail (Roche # 4693132001). Protein content in the supernatant was determined with BCA assay (Thermo #23225) and 25~50 µg of protein were subjected to SDS-PAGE and transferred to nitrocellulose membrane. Immunoblotting was performed with a mice anti-RAC2 antibody (Millipore 07-604-I), a rabbit anti-Vinculin antibody (Sigma V9131) and goat anti-mouse (Invitrogen 35518) or anti-rabbit (Thermo SA5-35571) secondary antibodies. The fluorescence intensity was measured using an Odyssey imaging system (LI-COR). The uncropped images are included in Supplemental Figure 2.

### Reporter assay

*miR-722* or vector control constructs were cloned into PCDNA3.1 as described (34). Human *RAC2* 3’UTR was amplified with One Step Ahead RT-PCR kit (Qiagen) from human mRNA using the following primers *RAC2* 3’UTR+: 5’-TAGGCGATCGCTCGAGGGGTTGCACCCCAGCGCT-3’, *RAC2* 3’UTR-: 5’-TTGCGGCCAGCGGCCGCTCTCCAAACTTGAATCAATAAATTT-3’ and inserted into a dual luciferase psiCHECK2 backbone (Promega). *RAC2* mutant 3’UTR constructs were generated using Infusion HD cloning kit (Clontech) with the following primers: *RAC2* mut+: 5’-GAGTTCCTCTACCCACGTTTTTTGAGTGTCTCAGAAGTGTGC-3’, *RAC2* mut–: 5’-TGGGTAGAGGAACTCTCTGG-3. Mutation was confirmed with Sanger sequencing. HEK-293T cells seeded at 2×10^5^/well in 12 well plates and transfected with the indicated *miR-722* overexpression and the RAC2 3’UTR reporter plasmids with Lipofectamine 3000 (Invitrogen). Cells were harvested and lysed at 48 h post transfection and luciferase activity was measured with Dual Glo-Luciferase reagent (Promega, PF-E2920) in a Synergy 2 (Biotek) plate reader. The renilla luciferase activity was normalized to the firefly luciferase, which was then normalized to the vector control.

### Transwell migration assay

HL-60 transwell assays were performed as described (46). Briefly, differentiated cells were resuspended in HBSS buffer and allowed to migration for 2 h towards fMLP (100 nM). Cells that migrated to the lower chamber were released with 0.5M EDTA and counted using a BD LSRFortessa flow cytometer with an acquisition time of 30 sec. The counts were normalized with the total numbers of cells added to each well the data was then gated for live cells and analyzed with Beckman Kaluza 2.1 software. The raw counts are provided in Supplementary Table 2.

### Chemotaxis assay

Differentiated HL-60 cells were loaded in pre-assembled ibidi chemotaxis µ-slides (ibidi # 80326) following the manufacture instruction and incubated at 37ºC for 30 minutes. 15 ul of 1000 nM fMLP was loaded into the right reservoir and cell migration was recorded every 60 sec for 120 minutes using LSM 710 (with Ziess EC Plan-NEOFLUAR 10X/0.3 objective) at 37ºC. Cells expressing Dendra2 with initial position on the left half of the channel were tracked with ImageJ plug-in MTrackJ as previously described (54). Velocity and chemotaxis index were quantified using the ibidi cell migration tool.

### Statistical Analysis

Statistical analysis was carried out using PRISM 6 (GraphPad). Mann–Whitney test (comparing two groups) and Kruskal–Wallis test (when comparing to single group) were used in this study as indicated in the figure legends. A representative experiment of at least three independent repeats were shown. For qPCR, each gene was normalized to the reference gene and compared with Mann–Whitney tests. The statistical significance of overlap between *miR-722* down-/up-regulated genes and *miR-722* target genes was calculated by using hypergeometric distribution.

## Acknowledgement

The authors would like to thank Dr. Daoguo Zhou (Purdue University) for providing the Odyssey imaging system (LI-COR). The work was supported by National Institutes of Health [R35GM119787 to DQ] and [P30CA023168 to Purdue Center for Cancer Research] for shared resources. Bioinformatics analysis was conducted by the Collaborative Core for Cancer Bioinformatics (C^3^B) shared by the Indiana University Simon Cancer Center [P30CA082709] and the Purdue University Center for Cancer Research with support from the Walther Cancer Foundation.

## Authorship

AH, JW and DQ designed research and wrote the manuscript. AH performed the experiments. AH, JW, RS, KB and SL analyzed the data. All authors read and approved the manuscript.

## Data availability statement

The RNA-seq raw data and processed data deposited to Gene Expression Omnibus (GEO) GSE126527. Plasmids are available on Addgene: pCDNA miR-722-Dendra (#97163), pcDNA3.1 Dendra2 (#103967), pSi-check2-hRac2 3′UTR (#97160), pSi-check2-hRac2 -mut 3′UTR (#97161), plix4.03-miR-722 (#97140), plix4.03-vector (#97141).

