## supplemental methods and figures for "Inducible overexpression of zebrafish *microRNA-722* suppresses chemotaxis of human neutrophil like cells"

**Supplementary Information**

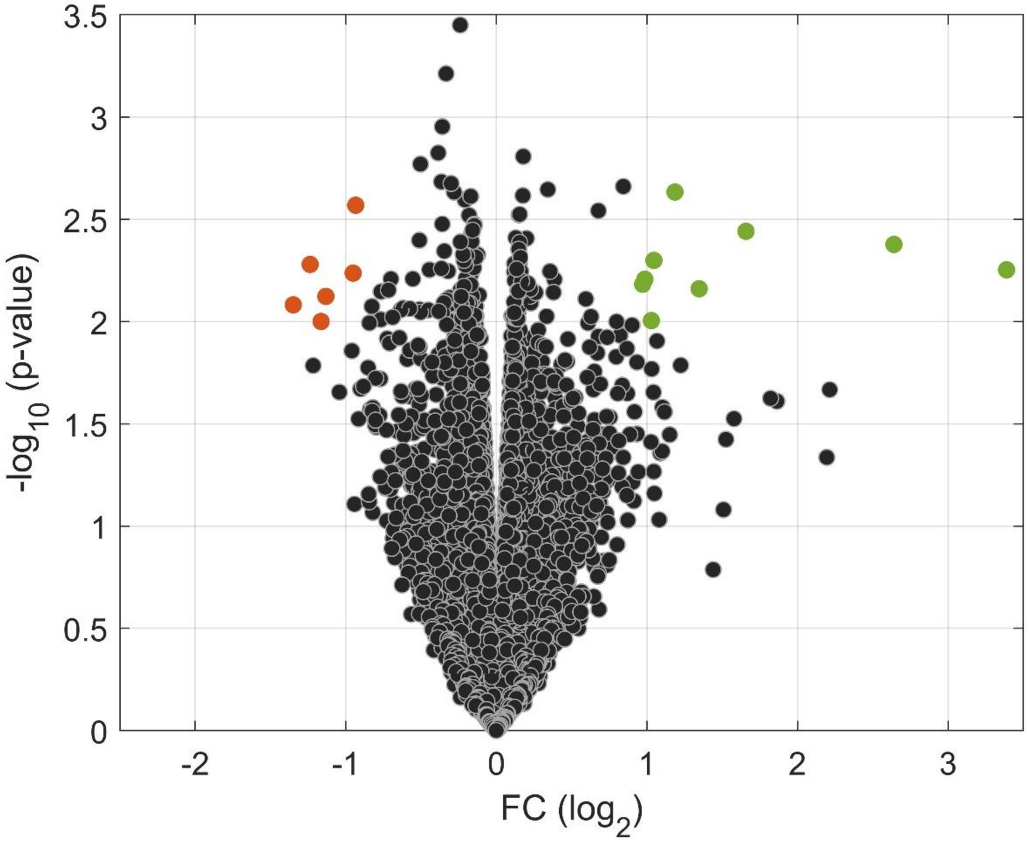


**Supplementary Figure 1. Inducible expression of Dendra2 in dHL-60 has minimal impact on the transcriptome.**
Volcano blot of DEGs with significant changes in expression upon doxycycline induced *Dendra2* expression. Orange: down regulated DEGs; green: up regulated DEGs.

**Supplementary Figure 2. Full length western blots.**

Immunoblot of RAC2 and the loading control Vinculin in vector or *miR-722* expressing dHL-60 cells ± DOX. The blot was cut in half and probed with two different primary antibodies.

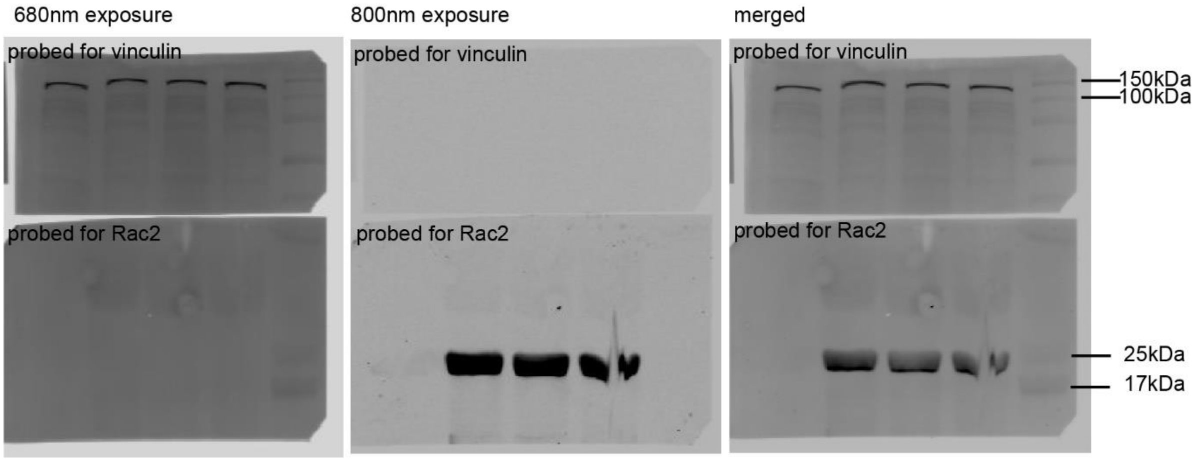

**Supplementary Table 1. Raw cell counts from FACS analysis of CD11b and Annexin V staining.**

| cells | CD11b | | Annexin V | |
|---|---|---|---|---|
| | number | %gated | number | %gated |
| vector-DOX | 10060 | 95.29 | 697 | 5.3 |
| vector+DOX | 10018 | 94.49 | 729 | 6.7 |
| miR-722-DOX | 10210 | 96.23 | 629 | 3.3 |
| miR-722+DOX | 10253 | 95.36 | 531 | 3 |

**Supplementary Table 2. Raw cell counts from FACS analysis of transwell migration.**

| | | Experiment 1 | | | | Experiment 2 | | | | Experiment 3 | | | |
|---|---|---|---|---|---|---|---|---|---|---|---|---|---|
| cell | treatment | migrated | loading | fraction | average | migrated | loading | fraction | average | migrated | loading | fraction | average |
| vector | Dox-FMLP+ (1) | 130 | 562 | 0.23132 | 0.24905 | 118 | 562 | 0.20996 | 0.21506 | 595 | 1869 | 0.31835 | 0.2788 |
| | Dox-FMLP+ (2) | 143 | 536 | 0.26679 | | 118 | 536 | 0.22015 | | 473 | 1977 | 0.23925 | |
| | Dox+FMLP-(1) | 8 | 503 | 0.0159 | 0.00965 | 4 | 503 | 0.00795 | 0.00823 | 93 | 1937 | 0.04801 | 0.0457 |
| | Dox+FMLP- (2) | 2 | 588 | 0.0034 | | 5 | 588 | 0.0085 | | 83 | 1913 | 0.04339 | |
| | Dox+FMLP+ (1) | 90 | 534 | 0.16854 | 0.17711 | 118 | 534 | 0.22097 | 0.17948 | 655 | 2159 | 0.30338 | 0.30978 |
| | Dox+FMLP+ (2) | 109 | 587 | 0.18569 | | 81 | 587 | 0.13799 | | 596 | 1885 | 0.31618 | |
| mir722 | Dox-FMLP+ (1) | 147 | 1363 | 0.10785 | 0.11534 | 131 | 1363 | 0.09611 | 0.10683 | 612 | 2254 | 0.27152 | 0.32883 |
| | Dox-FMLP+ (2) | 163 | 1327 | 0.12283 | | 156 | 1327 | 0.11756 | | 763 | 1976 | 0.38613 | |
| | Dox+FMLP-(1) | 3 | 1595 | 0.00188 | 0.00094 | 2 | 1595 | 0.00125 | 0.00115 | 44 | 1895 | 0.02322 | 0.02523 |
| | Dox+FMLP- (2) | 0 | 1901 | 0 | | 2 | 1901 | 0.00105 | | 57 | 2093 | 0.02723 | |
| | Dox+FMLP+ (1) | 4 | 1830 | 0.00219 | 0.0019 | 2 | 1830 | 0.00109 | 0.00135 | 109 | 2403 | 0.04536 | 0.05111 |
| | Dox+FMLP+ (2) | 3 | 1860 | 0.00161 | | 3 | 1860 | 0.00161 | | 112 | 1970 | 0.05685 | |

**Supplementary Dataset 1. List of genes with significant expressional change upon induced *miR-722* expression.**

**Supplementary Dataset 2. List of genes with significant expressional change upon induced expression of the reporter gene alone.**

**Supplementary Dataset 3. Pathway analysis of differentially expressed genes in the *miR-722* line upon doxycycline mediated induction.**

**Supplementary Movie 1. Chemotaxis of HL-60 upon *miR-722* overexpression.**

Representative movies of dHL-60 cells expressing either vector (left) or *miR-722* with (right) and without (middle) doxycycline toward fMLP. Scale bar: 100 µm.

**Supplementary Movie 2. Tracks of HL-60 chemotaxis upon *miR-722* overexpression.**

Tracks of Movie S1. All tracks are assigned to start at (0,0). Red crosses indicate the center of mass at each time point.
